# Speaker gaze increases information coupling between infant and adult brains

**DOI:** 10.1101/108878

**Authors:** Victoria Leong, Elizabeth Byrne, Kaili Clackson, Stanimira Georgieva, Sarah Lam, Sam Wass

**Affiliations:** Department of Psychology, University of Cambridge; Nanyang Technological University, Singapore; MRC Cognition & Brain Sciences Unit, Cambridge; University of East London

**Keywords:** Neural synchronisation, dyadic interaction, mutual gaze, intention

## Abstract

When infants and adults communicate, they exchange social signals of availability and communicative intention such as eye gaze. Previous research indicates that when communication is successful, close temporal dependencies arise between adult speakers’ and listeners’ neural activity. However, it is not known whether similar neural contingencies exist within adult-*infant* dyads. Here, we used dual-electroencephalography to assess whether direct gaze increases neural coupling between adults and infants during screen-based and live interactions. In Experiment 1 (N=17), infants viewed videos of an adult who was singing nursery rhymes with (a) ***Direct gaze*** (looking forward); (b) ***Indirect gaze*** (head and eyes averted by 20°); or (c) ***Direct-Oblique gaze*** (head averted but eyes orientated forward). In Experiment 2 (N=19), infants viewed the same adult in a live context, singing with Direct or Indirect gaze. Gaze-related changes in adult-infant neural network connectivity were measured using Partial Directed Coherence. Across both experiments, the adult had a significant (Granger)-causal influence on infants’ neural activity, which was stronger during Direct and Direct-Oblique gaze relative to Indirect gaze. During live interactions, infants also influenced the adult more during Direct than Indirect gaze. Further, infants vocalised more frequently during live Direct gaze, and individual infants who vocalized longer also elicited stronger synchronisation from the adult. These results demonstrate that direct gaze strengthens bi-directional adult-infant neural connectivity during communication. Thus, ostensive social signals could act to bring brains into mutual temporal alignment, creating a joint-networked state that is structured to facilitate information transfer during early communication and learning.

## INTRODUCTION

### Gaze in early development

Temporally contingent social interactions between adults and infants play a vital role in supporting early learning across multiple domains of language, cognition and socio-emotional development [1,2]. Infants rely heavily on the temporal dynamics of facial cues such as eye contact and gaze direction to infer intention, meaning and causality [3-5], which is unsurprising given that infants’ early visual experience is heavily composed of faces [6]. Of all cues, direct gaze is thought to be one of the most salient ostensive signals in human communication for conveying communicative intent [4]. Gaze also acts to release and reinforce infants’ own social responses such as smiling and vocalisation [7,8]. From birth, infants prefer to look at pictures of faces with direct gaze over averted gaze [9]. By 4 months, direct gaze elicits a larger amplitude in the face-sensitive N170 event-related potential (ERP) relative to averted gaze [10], which suggests that gaze also enhances infants’ neural processing of face-related information.

### Social synchronisation through gaze in communication

According to the social brain hypothesis, human brains have fundamentally evolved for group living [11]. Social connectedness is created when group members act jointly (e.g. synchronously) or contingently (e.g. turn-taking) with each other [12]. Even infants show synchronisation with their adult caregivers, and adult-infant temporal contingencies have long been observed in behavioural and physiological domains. For example, patterns of temporally synchronous activity between parent and child during social interaction have been noted for gaze [13], vocalisations [14], affect [15], autonomic arousal [16,17], and hormones [18]. The synchronisation of gaze (through mutual gaze and gaze-following) is thought to foster social connectedness between infants and adults [19]. Previous research has also suggested that infants, like adults [20], show *neural* synchronisation (or phase-locking) of cortical oscillatory activity to temporal structures in auditory signals [21]. However, adult-infant behavioural and physiological synchronisation is typically observed over much slower timescales (e.g. minutes or seconds) than neural synchronisation (tens or hundreds of milliseconds). Thus, it remains to be seen whether neural synchronisation also develops between infants and adults during social interaction, and if/how such neural coupling is related to social synchronising signals like gaze.

Recently, researchers have begun to examine the neural mechanisms which support the contingency (temporal dependency) of one partner’s neural activity with respect to the other during social interactions (see [22,23] for reviews). This work has revealed that during verbal communication (especially face-to-face communication which permits mutual gaze), adult speaker-listener pairs develop synchronous patterns of activity between brain regions such as the inferior frontal gyrus, prefrontal and parietal cortices [24,25]. Further, the strength of speaker-listener neural synchronisation predicts communication success [26]. Thus, in adults, effective communication involves the mutual alignment of brain activity, as well as the temporal alignment of behaviour (e.g. conversational turn-taking and mutual gaze). Yet to our knowledge, no previous research has yet investigated whether infants’ neural activity also shows contingency on an adult partner’s neural activity, and whether gaze acts as a neural synchronisation cue during adult-infant communication.

### Gaze-cueing of interpersonal neural synchronisation

Here, we assessed whether the temporal dependency (synchronisation) between adult and infant neural signals differed between Direct and Indirect gaze. Two experiments were performed to assess gaze-cueing of interpersonal synchronisation in video and live modalities respectively. In Experiment 1, infants watched a pre-recorded video of an experimenter singing nursery rhymes. Patterns of temporal dependency were assessed between infants’ neural activity recorded ‘live’ and adult’s pre-recorded neural activity (see Figure 1). We manipulated the adult speaker’s gaze to either be Direct to the infant, Indirect (head averted at a 20° angle), or Direct-Oblique (head averted but eyes toward the infant). The Direct-Oblique condition was included to control for the side view of the face that was presented during Indirect gaze, and to preclude the possibility that infants were responding to superficial visual differences between stimuli. In Experiment 2, which used an entirely separate cohort, infants listened live to an adult reciting nursery rhymes whilst she presented Direct or Indirect gaze to the infant. Partial directed coherence [27], a statistical measure of Granger causality [28], was used to measure gaze-related changes in interpersonal neural synchronisation within the adult-infant dyadic social network.

**Figure 1.**
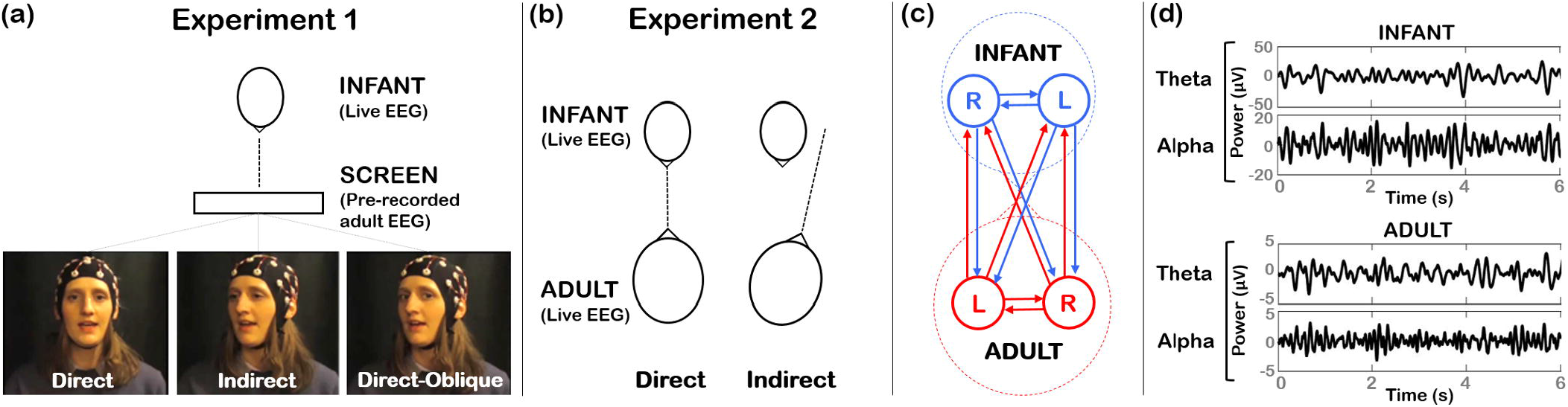
*Illustration of experimental protocols and connectivity analysis (a) In Expt 1 infants viewed a video screen showing an experimenter reciting nursery rhymes. Three gaze conditions were presented interleaved: Direct, Indirect (head averted by 20°), and Direct-oblique (head averted by 20°, direct gaze). The infant’s live EEG was compared with the adult’s pre-recorded EEG. (b) In Expt 2, infant and adult sat opposite each other. Direct and Indirect gaze (head averted by 20°) conditions were presented. (c) The adult-infant network comprised left and right electrodes each from the infant and adult. Interpersonal neural connectivity was assessed across all pairwise connections between electrodes using partial directed coherence. (d) Examples of infant and adult EEG data, which was analysed within Theta (3-6 Hz) and Alpha (6-9 Hz) bands.*

### Predictions

In terms of affect and physiological changes, research has shown that the influence of infants and parents on one another is bi-directional [29,30]. Accordingly, we predicted that: i) significant neural coupling would exist between adults and infants during social interaction; ii) Direct (and Direct-Oblique) gaze would both be associated with higher interpersonal neural connectivity than Indirect gaze; and iii) in Experiment 1 (Video), only unidirectional (adult-to-infant [A→I]) coupling would be observed, but in Experiment 2 (Live), bi-directional (adult-to-infant [A→I] and infant-to-adult [I→A]) coupling would be observed. Further, as temporally contingent social interactions with adults are known to facilitate infants’ own vocalisations [8,31], we predicted that infants’ vocalisation efforts would be greater during Direct than Indirect gaze.

## RESULTS

### Gaze Modulation of Interpersonal Neural Connectivity

General Partial Directed Coherence (GPDC) measures the degree of influence that each electrode channel *directly* has on every other electrode channel in the network [27]. Here, GPDC values were computed for real and surrogate (shuffled) data, for all non-self channel pairs (connections), for each participant dyad, for each gaze condition, and in Theta and Alpha EEG bands (see Figure 1c & 1d). In the subsequent network diagrams (Figures 2 & 3), only connections whose GPDC values significantly exceeded their surrogate threshold are plotted. A breakdown of GPDC values for each neural connection is provided in SI Appendix Section 1 (Tables S1 & S2). Here we focus our analysis on *mean* adult-to-infant (A→I) and infant-to-adult (I→A) connectivity.

**Figure 2.**
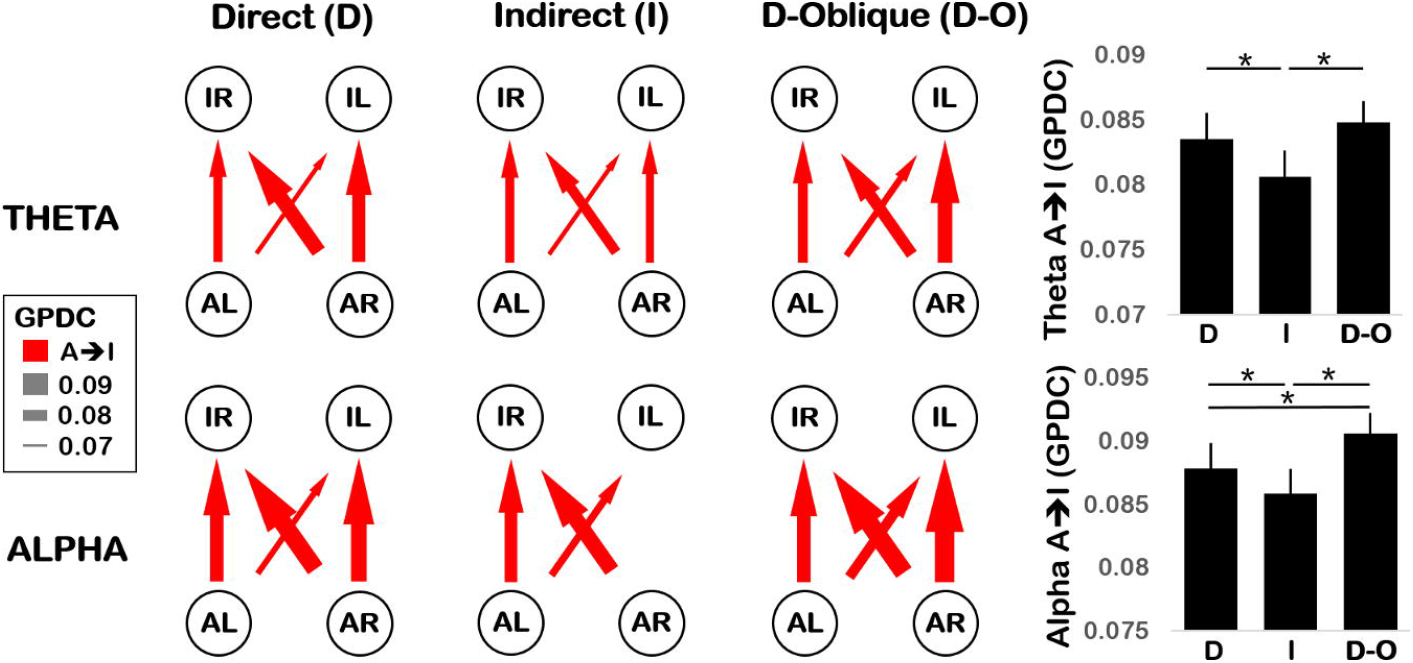
*(Left) Network depiction of Expt 1 Theta (3-6 Hz, top row) and Alpha (6-9 Hz, bottom row) connectivity, plotting GPDC values for Direct (left column), Indirect (middle column) and Direct-Oblique gaze (right column) conditions. Nodes represent C3 (L) and C4 (R) electrodes for adult (A) and infant (I). Arrows indicate the direction and strength of connectivity (higher GPDC value = thicker arrow). Connections that do not significantly exceed the surrogate threshold are excluded. (Right) Grand mean GPDC values averaged across all adult-to-infant (A→ I) connections for Theta (top) and Alpha (bottom) in Direct (D), Indirect (I) and Direct-Oblique (D-O) gaze conditions. Error bars show the standard error of the mean. *p<.05*

**Figure 3.**
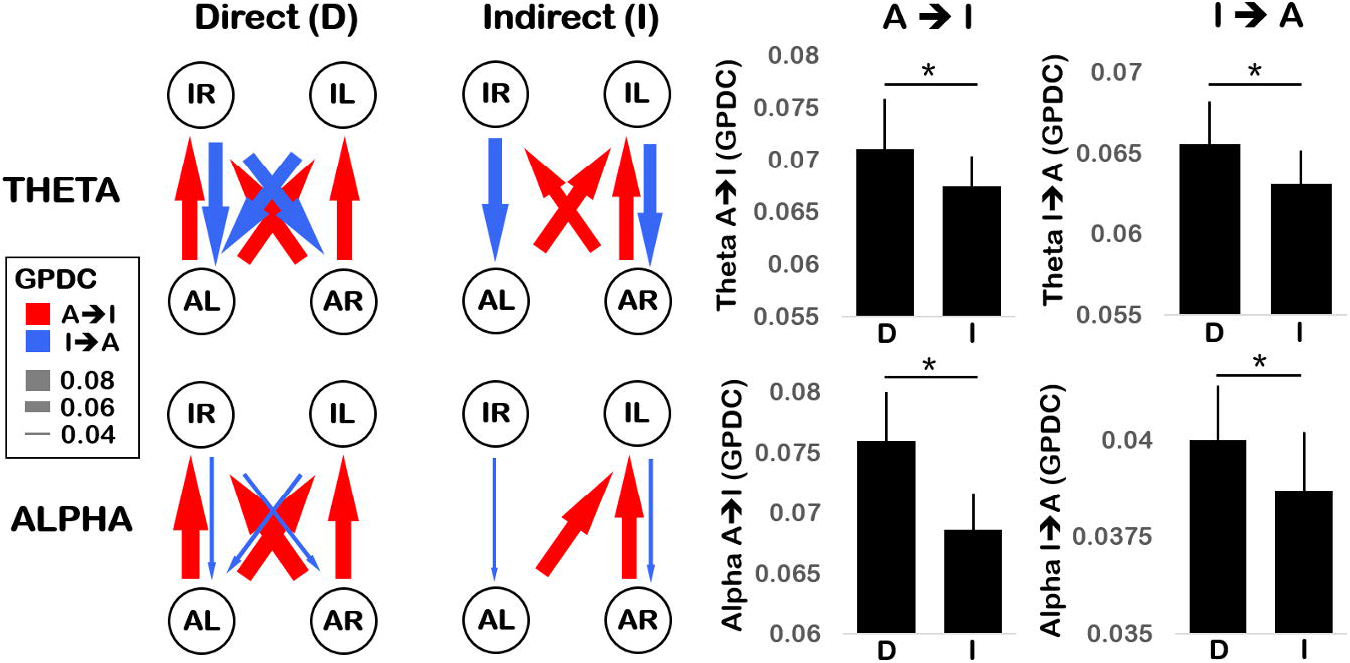
*(Left) Network depiction of Expt 2 Theta (3-6 Hz, top row) and Alpha (6-9 Hz, bottom row) connectivity, plotting GPDC values for Direct (left column) and Indirect (right column) gaze conditions. Nodes represent C3 (L) and C4 (R) electrodes for adult (A) and infant (I). Arrows indicate the direction and strength of connectivity (higher GPDC value = thicker arrow). Connections that do not significantly exceed the surrogate threshold are excluded. (Right) Grand mean GPDC values averaged across all adult-to-infant (A→ I, left column) and infant-to-adult (I → A, right column) connections for Theta (top row) and Alpha (bottom row) in Direct (D) and Indirect (I) gaze conditions. Error bars show the standard error of the mean. *p<.05*

#### Experiment 1: Video

Only *uni-directional* A→I connectivity was observed in Experiment 1, *no* significant I→A connectivity was detected (see Figure 2). This confirmed the validity of the GPDC measure as infants could not have affected the adult’s pre-recorded neural activity. Dunnett’s tests revealed that, as predicted, A→I connectivity was (1) significantly stronger for ***Direct > Indirect*** gaze in both Theta and Alpha bands (p<.01, p<.05 respectively, one-tailed); and (2) significantly stronger for ***Direct-Oblique > Indirect*** gaze in both Theta and Alpha bands (p<.0001 for both, one-tailed). However, whilst connectivity in the Direct and Direct-Oblique conditions was not significantly different in the Theta band (p=.30) as predicted, for the Alpha band a significant difference between these conditions was observed (***Direct-Oblique > Direct***, p<.01).

#### Experiment 2: Live

During the live experiment, *bi-directional* connectivity was observed with significant A→I as well as I→A influences (see Figure 3).

*Adult-to-infant (A→I) connectivity.* Consistent with Experiment 1, Dunnett’s tests revealed that A→I was significantly stronger for ***Direct > Indirect*** gaze in both Theta and Alpha bands (p<.05 and p<.0001 respectively, one-tailed).

*Infant-to-adult (I→A) connectivity.* Regarding infants’ influence on the adult, Dunnett’s tests revealed that, likewise, I→A was also significantly stronger for ***Direct > Indirect*** gaze in both Theta and Alpha bands (p<.01 and p<.05 respectively, one-tailed).

### Infant Vocalisation Analysis

For Experiment 1 (video), there was no difference in the number of infant vocalisations (summed over all categories) between gaze conditions (means: Direct = 8.2 per infant, Indirect = 7.4, Direct-Oblique = 7.1; F(2, 32) = .29, p=.75, η^2^p = .02). There was also no difference in the duration of vocalisations across gaze conditions (means : Direct = 0.69s per utterance, Indirect = 0.82s; Direct-Oblique = 0.70s; (F(2, 24) = .37, p=.70, η^2^p = .03). However, for Experiment 2 (live), we observed a significantly higher number of vocalisations during Direct gaze (mean 6.3 per infant) than Indirect gaze (mean 5.0 per infant; t(18) = 2.41, p<.05), but no difference in the duration of vocalisations (mean : Direct = 0.80s per utterance, Indirect = 0.85s; t(15) = -.79, p = .44).

Further, during Experiment 2 (live), individual differences in infants’ vocalisation duration were significantly associated with their I→A GPDC values (r=.67, p<.05, Benjamini-Hochberg FDR corrected [32]), see Figure 4. However, this correlation only emerged during Direct gaze, and was absent for Indirect gaze (r=.07, p=.78). Therefore, infants who produced longer vocalisations also influenced the adult more strongly – but only when she offered Direct gaze. SI Appendix Section 2 provides further analyses of infants’ vocalisations.

**Figure 4.**
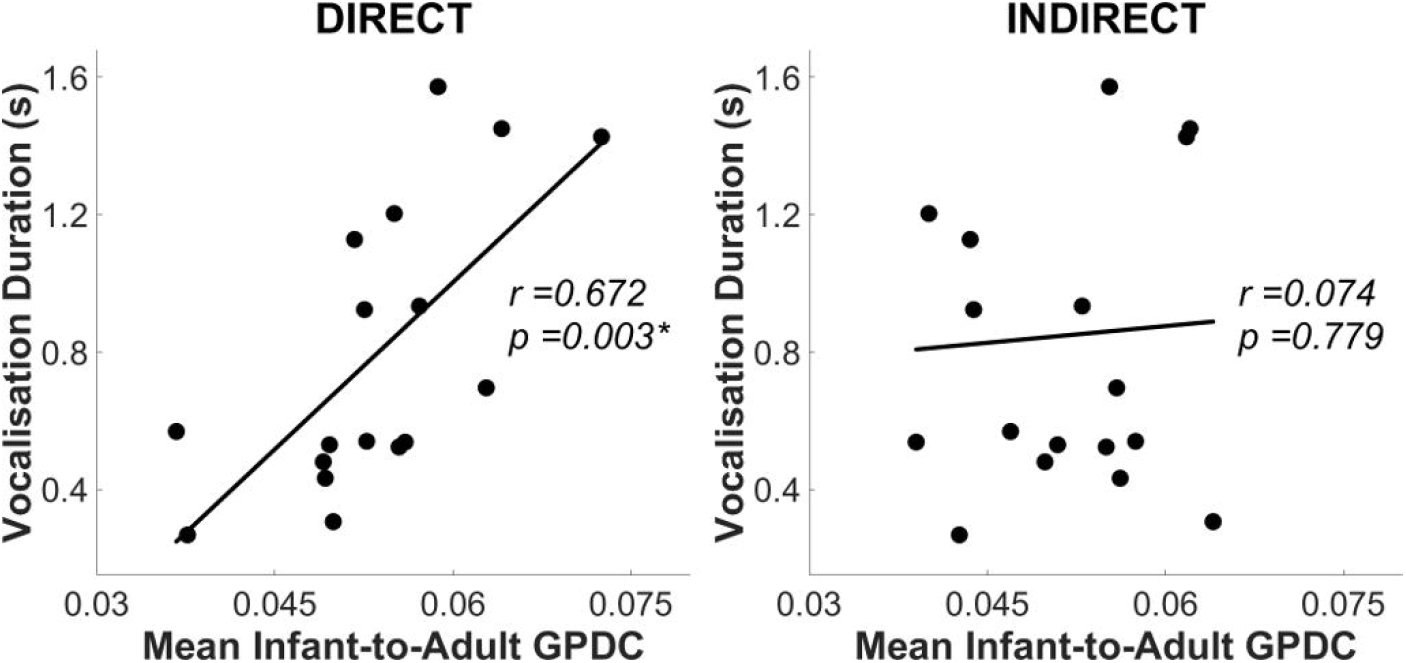
*Scatterplots showing the correlation between (N=19) individual infants’ mean Infant-to-Adult GPDC values (averaged across Theta and Alpha bands, x-axis), and their vocalisation duration (y-axis) in Experiment 2. Left and right plots show Direct and Indirect gaze conditions respectively. *p<.05 (BH FDR corrected)*

## DISCUSSION

Temporally contingent social interactions between adults and infants scaffold early learning and development. Here, we tested the hypothesis that gaze acts as an interpersonal neural synchronisation cue between dyadic (adult-infant) partners. Two experiments were performed to assess the effect of Direct speaker gaze on interpersonal synchronisation using video (Experiment 1) and live (Experiment 2) modalities. Across both experiments, significant neural coupling between infants and adults was observed during social interaction, relative to rigorous control analyses that accounted for non-specific neural coupling. Adult-infant neural coupling was observed consistently across Video and Live presentation formats, using two separate cohorts of infants. Further, during uni-directional interactions in Experiment 1 (i.e. infants watching a pre-recorded adult speaker), the adult had a significant influence on infants’ neural activity, but (as expected) infants had no influence on the adult’s neural activity. Conversely, during live (bi-directional) social interactions (Experiment 2), there were significant and bi-directional patterns of influence between adult and infant.

Across both experiments, we consistently observed that Direct gaze produced higher interpersonal neural synchronisation than Indirect gaze in both Theta and Alpha frequency bands. Further, in Experiment 2 (live), the synchronizing effect of gaze was observed bi-directionally: during Direct gaze, the adult had a stronger influence on the infant, and the infant also had a stronger influence on the adult. This gaze-related increase in synchronisation was not due to power differences in the EEG spectra, nor was it a meta-phenomenon of changes in basic sensory processing of the speech signal (which remained unchanged across gaze conditions). In Experiment 1, we further showed that the gaze effect was not driven by superficial visual differences in the stimuli, since Direct-Oblique stimuli were visually-similar to Indirect stimuli but produced greater synchronisation. It was also not the case that infants were more inattentive during Indirect gaze, as infants looked just as long at Indirect and Direct-Oblique stimuli in Experiment 1, and at Indirect and Direct stimuli in Experiment 2. Therefore, the increased interpersonal neural synchronisation produced by direct gaze appears to reflect stronger mutual oscillatory phase-alignment between adult and infant.

### A mechanism for interpersonal neural synchronisation

One mechanism that might mediate this effect is mutual phase-resetting in response to salient social signals. The *phase* of cortical oscillations (the neural feature used in GPDC computations) reflects the excitability of underlying neuronal populations to incoming sensory stimulation [33]. Sensory information arriving during high receptivity periods is more likely to be encoded than information arriving during low receptivity periods. Consequently, neuronal oscillations have been proposed to be a mechanism for temporal sampling of the environment [20]. Specifically, salient events are thought to *reset* the phase of on-going neuronal oscillations to match the temporal structure of these events and optimise their encoding [33]. Consequently, interpersonal neural synchronisation could increase within a dyad during the course of social interaction because each partner is continuously producing salient social signals (such as gaze, gestures, or vocalisations) that act as synchronisation triggers to reset the phase of their partner’s on-going oscillations. As a result, infants’ most receptive periods become well-aligned to adults’ speech temporal patterns (e.g. prosodic stress and syllable patterns [34]), optimising communicative efficiency. This mechanism could also allow slow-varying *behavioural* synchronisation signals (like gaze) to hierarchically control fast-varying *neural* synchronisation between partners [33].

### Direct gaze supports communication through synchronisation

Our findings suggest that direct gaze from the adult may reset the phase of infants’ oscillations to align with that of the adults’, thereby increasing mutual synchronisation (i.e. stronger A → I connectivity). One aspect of our results was, however, unpredicted. In Experiment 1, we had predicted an equal effect for Direct and Direct-Oblique gaze, yet we found that Alpha neural synchrony was *higher* for Direct-Oblique than Direct gaze. One possible explanation for this is that infants are less frequently exposed to direct eye contact when the speaker’s head is averted, which could therefore present greater novelty. However, infants did not look for longer at the speaker during the Direct-Oblique condition relative to the Direct gaze condition, which is inconsistent with this explanation. A second potential explanation is that the Direct-Oblique condition provided a stronger *intentional* ostensive cue because the speaker’s gaze was intentionally forward while her face and body were averted. This predicts that social cues which are perceived as the most intentional will produce the strongest increases in interpersonal connectivity. Further, since phase-resetting optimises information transfer between dyadic partners [33], stronger intentional signals could produce more effective phase-resetting, which would increase the potential for mutual communication and learning within the dyad. Future work should investigate this hypothesis in more detail.

As observed in previous studies [8], we also found that infants vocalised more frequently toward the adult during live Direct gaze (when interpersonal synchronisation was higher) than Indirect gaze. Further, individual infants who vocalized for longer under live Direct gaze also had stronger neural connectivity with their adult partner (i.e. stronger I → A connectivity), even during segments when no vocalisations were occurring. One possible reason for this could be that infants’ vocalisations (which were communicative signals to the adult and could potentially trigger phase-resetting), acted as a social feedback mechanism to positively reinforce and sustain dyadic synchronicity [8,31,35].

Our present findings may offer the potential for integrating three separate strands of research into early learning: first, research that has pointed to the importance of eye gaze as an ostensive cue during learning [3]; second, research into the importance of contingent social feedback which is thought to energise early learning [31]; third, research into the role of bi-directional parent-child synchrony in structuring and scaffolding learning experiences [36]. Phase-resetting due to synchronisation triggers that are more prevalent during mutual than indirect gaze may, potentially, offer the means for providing *contingent* feedback (in which the child responds to the parent, and *vice versa*) within the framework of the periodic oscillatory activity that structures and scaffolds early learning [36]. Over longer time frames, infants’ neural synchrony with adults may also offer an implicit mechanism for learning adult-like response patterns via entrainment.

### Limitations and Conclusion

Our results converge with previous dual-fNIRS studies [24,37] where greater frontal neural synchronisation between adults was observed during eye-contact. However, one limitation of the current work is that due to the adult’s speech production artifacts, only two EEG channels, C3 and C4, could be analysed from each individual. Thus, unlike the fNIRS studies, we were unable to make inferences about the potential neural sources of these effects. A second limitation of the current work is that, by excluding a large proportion of infants’ ‘active’ data by technical necessity, this could present a selective view of the neural dynamics underlying adult-infant engagement. Nonetheless, the current data are still valuable in providing insight into adult-infant neural coupling during social communication.

The current study demonstrates that adults and infants show significant mutual neural coupling during social interactions, and that direct gaze strengthens adult-infant neural connectivity in both directions during communication. Further, live gaze appeared to stimulate infants’ own communicative efforts which could help to reinforce dyadic synchronisation. Thus, gaze and speech act as cues for interpersonal synchronisation. The contingent exchange of these social signals acts to bring adults’ and infants’ brains into temporal alignment, creating a joint-networked state that is structured to optimise information transfer during communication and learning.

## METHODS

### Participants

Experiments 1 and 2 involved separate infant cohorts. Expt 1: Nineteen infants (13M, 6F), median age 8.2 m (SE : 0.26 m). Expt 2: Twenty-nine infants (15M, 14F), median age 8.3 m (SE : 0.44 m). Infants’ mothers were native English speakers and all infants had no neurological problems as assessed by maternal report. The same female adult experimenter participated in both experiments with all infants. The study received ethical approval from the Cambridge Psychology Research Ethics Committee. Parents provided written informed consent on behalf of their infants.

### Materials

For both experiments, seven familiar nursery rhymes were used as sung stimuli (see SI Appendix Section 3). Sung nursery rhymes were used because these are integral to play and caretaking routines with infants, such as during feeding and putting to sleep [38]. Infants are equally or more behaviourally responsive to sung as compared to spoken language [39], thus sung speech is likely to evoke a robust neural response from infants. In Experiment 1, pre-recorded video stimuli were used with mean pitch, pitch variability, duration and loudness matched across gaze conditions (SI Appendix Table S5). For Experiment 2 (live), the experimenter was recorded during each session to ensure acoustic consistency across gaze conditions (SI Appendix Table S6). Paired t-tests indicated no significant differences between conditions for all acoustic parameters. The experimenter was instructed to maintain a neutral facial expression across all gaze conditions, varying only her gaze direction.

### Protocol

#### Experiment 1

Infants sat upright in a high chair 70 cm from a display monitor (90 cm W x 60 cm H), showing a life-sized image of a female experimenter’s head against a black background. Each nursery rhyme was presented in three gaze conditions (see Figure 1): Direct, Indirect (head averted by 20°) and Direct-Oblique (head averted by 20°, but direct gaze). The Direct-Oblique condition was included to control for the side view of the face that was presented during Indirect gaze. During stimulus recording, the experimenter gaze-fixated on a life-sized picture of an infant to standardise her visual input across conditions. Each nursery rhyme was presented six times (twice per gaze condition, order counterbalanced).

#### Experiment 2

Infants sat upright in a high chair facing the female experimenter at a distance of 70 cm. Each nursery rhyme was presented in two gaze conditions. In the Direct condition the experimenter looked directly at the infant while singing; in the Indirect condition she fixated at a target 20° to the left or right side of the infant (see Figure 1, and SI Appendix Section 4 for the experimenter’s view). Each nursery rhyme was presented four times (twice Direct, twice Indirect, order counterbalanced).

### EEG acquisition

In Experiment 1, EEG was recorded *separately* from infants (during testing) and from the female adult experimenter (during stimulus recording) from 32 electrodes according to the International 10–20 placement system. In Experiment 2, EEG was recorded *simultaneously* from the infant and the adult experimenter from two central electrodes (C3 and C4), referenced to the vertex (Cz). Further details of EEG acquisition are given in SI Appendix Section 5.

### EEG artifact rejection and pre-processing

To ensure that the analysed EEG data reflected only attentive and movement-free neural activity, a two-stage artifact rejection procedure was applied. First, session videos were manually-reviewed to select only periods when infants were still and looking directly at the experimenter. Next, manual artifact rejection was performed to further exclude segments where the EEG amplitude exceeded +100 μV. Full descriptions of the artifact rejection procedures and inclusion rates following artifact rejection are given in SI Appendix Section 6. Data were then downsampled to 200 Hz, low-pass filtered <45 Hz to suppress electrical line noise, and segmented into 1.0s epochs for connectivity analysis.

### EEG analyses: Speech artifacts, power spectrum and GPDC network connectivity

Speech production artifacts were present in the EEG signal of the adult speaker. To assess the topography and spectral profile of these artifacts, we compared the adult’s EEG during speech production relative to resting state (see SI Appendix Section 7). Despite rigorous analyses we were able to identify no evidence of EEG signal distortion by speech artifacts in the central region (e.g. C3/C4) in Theta and Alpha bands, although evidence of artifacts at other frequency bands and for more peripheral electrode positions was clearly present. Therefore, to avoid spurious results arising from speech artifacts, the connectivity analysis used only Theta and Alpha bands for C3 and C4 electrodes for both adult and infant. To confirm the representativeness of this region of analysis for the infant, we assessed infants’ whole-head (32-channel) connectivity to adults’ C3 and C4 electrodes (see Figure 5 and SI Appendix Section 12). Across gaze conditions, the strongest connectivity between infant and adult was topographically observed over infants’ central and posterior regions (including C3 and C4) for both Theta and Alpha bands. Therefore, C3 and C4 were indeed representative regions of analysis for the infant.

**Figure 5.**
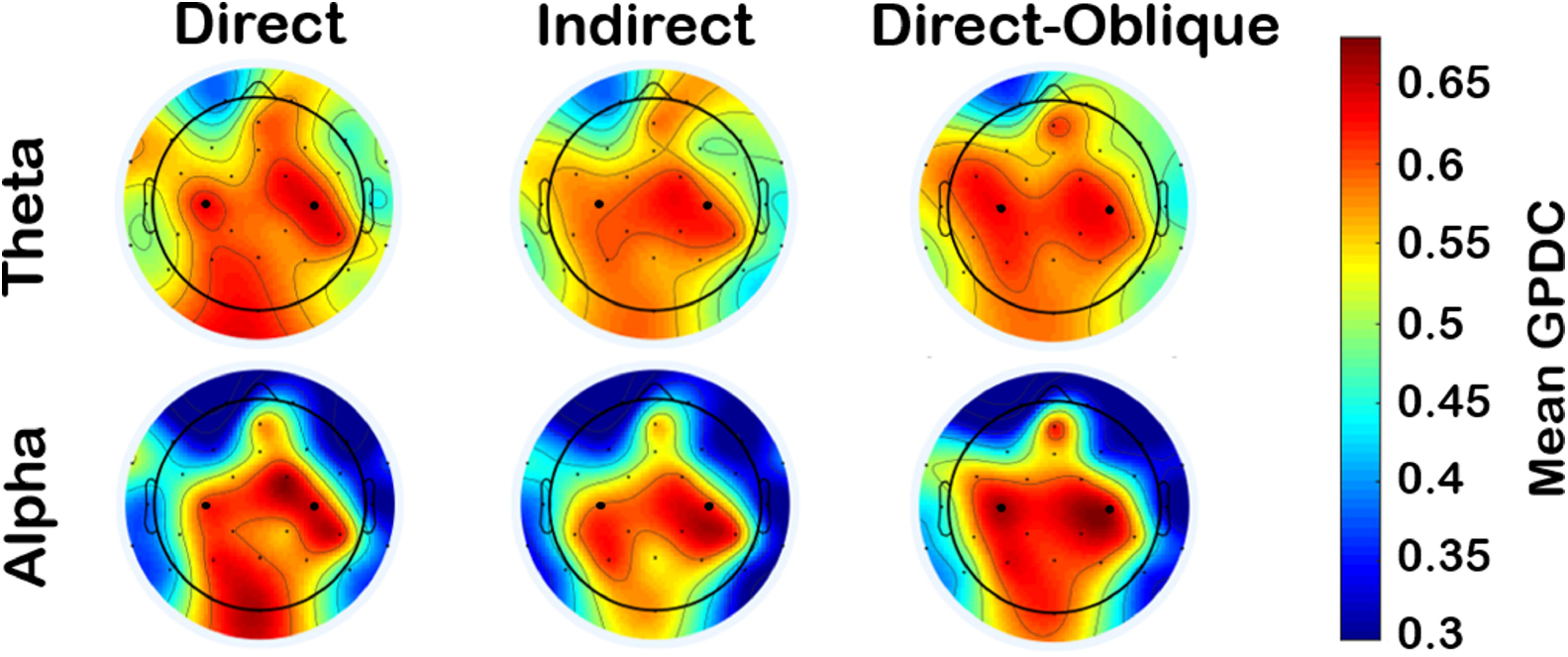
*Experiment 1 infant scalp topography of the mean adult (C3/C4)-to-infant GPDC values for Direct gaze (left column), Indirect gaze (middle column) and Direct-Oblique gaze (right column) conditions, for Theta (top row) and Alpha (bottom row) frequency bands. Electrodes C3 and C4 are enlarged for ease of reference. For each subplot, a top-down view of the scalp is shown where left/right map congruently to left/right sides of the infant’s head respectively.*

A detailed description of EEG analysis methods is given in SI Appendix Sections 8 to 9. Briefly, first the EEG power spectra of infant and adult signals were assessed for each experimental condition to confirm that the gaze manipulation did not generate any detectable power changes that might systematically bias the connectivity analysis. Second, to assess network connectivity in each gaze condition, Generalised Partial Directed Coherence (GPDC) was computed - a directional causal measure of direct information flow between channels in a network [27]. GPDC measures the degree of influence that channel i *directly* has on channel j with respect to the total influence of i on all channels in the network. Here, each electrode (IL, IR, AL, AR) was one channel (see Figure 1c).

### Control analyses

The first control analysis established a threshold for non-specific connectivity between brains that was unrelated to the experimental task (see SI Appendix Section 10). A surrogate dataset was generated for each participant pair where the fine-grained temporal correspondence between adult and infant neural signals was disrupted by randomly pairing adult and infant epochs from different timepoints within the same experimental session (i.e. shuffling). An identical connectivity analysis was then performed on this surrogate dataset. For each participant pair, neural connection and frequency band, a threshold value was computed by taking the average surrogate value across all gaze conditions. Paired t-tests (BH-corrected at p<.05 [32], one-tailed) were then used to assess whether the real data significantly exceeded their respective threshold values.

The second control analysis examined basic sensory processing of the speech stimulus which could indirectly affect adult-infant neural coupling. Entrainment (oscillatory phase-locking) between the EEG signal and the speech amplitude envelope was measured in each gaze condition. As described in SI Appendix Section 11, no significant differences in neural entrainment to the speech signal between gaze conditions were found in either experiment.

### Statistical analysis of gaze effects on interpersonal GPDC connectivity

We hypothesised that interpersonal neural connectivity would be higher during Direct (and Direct-Oblique) gaze than Indirect gaze (i.e. Direct=Direct-Oblique > Indirect). We also wished to assess whether the adult’s influence on the infant (i.e. adult-to-infant [A→I] GPDC) and the infant’s influence on the adult (i.e. infant-to-adult [I→A] GPDC) would show the same pattern of gaze modulation. As previous work with infants has not found hemispheric differences for gaze effects [9], interhemispheric connectivity patterns were not explored further. Accordingly, the four interhemispheric connections (L/R → L/R) were collapsed into one average each for A→I and I→A directional influences. These two directional indices were computed for each gaze condition, for Theta and Alpha bands. For Expt 1, only A→I connections were analysed as all I→A connections were not significantly above threshold (this was expected as the adult’s EEG was pre-recorded).

The effects of gaze on A→I and I→A connectivity were assessed using two statistical approaches. First, to assess *overall patterns* and interactions, Repeated Measures (RM) ANOVAs were performed, taking Frequency and Gaze condition as within-subjects factors. Second, to assess *specific contrasts* between pairs of gaze conditions at each frequency, Dunnett’s multiple range t-tests [40] were conducted, which independently control for the familywise error rate. For Theta and Alpha bands, the following pairwise tests were performed for Expt 1: ***[1] Direct > Indirect***; ***[2] Direct-Oblique > Indirect***; and ***[3] Direct = Direct-Oblique***. For Expt 2, only the ***Direct > Indirect*** test was performed. Dunnett’s test results are reported in the main manuscript, and ANOVA results are provided in SI Appendix Section 13. Separate analyses were also performed to examine infants’ looking times (SI Appendix Section 14) and the effects of infant age on neural connectivity (SI Appendix Section 15). Finally, a permutation analysis was performed (SI Appendix Section 16) to assess the internal reliability of the gaze findings, both within and across experiments. All statistical tests were two-tailed unless there were *a-priori* directional hypotheses (i.e. Dunnett’s test for Direct/Direct-Oblique > Indirect; Data > Surrogate threshold), for which one-tailed tests were used.

### Infant Vocalisations

Infants’ vocalisations were coded from session videos according to Oller’s [41] infraphonological acoustic classification system (see SI Appendix Section 2). Each infant’s (a) number and (b) duration of vocalisations was computed during each gaze condition. To explore the relationship between neural coupling and infants’ communicative attempts, vocalisation indices were correlated with A→I and I→A GPDC values for both experiments. Of note, the connectivity analyses only included segments of EEG data when no vocalisations were occurring.

## ACKNOWLEDGEMENTS

This research was funded by an ESRC Transforming Social Sciences grant (ES/N006461/1) to VL and SW, a Lucy Cavendish College Junior Research Fellowship and a Nanyang Technological University start-up grant M4081585.SS0 to VL, and a British Academy Post-Doctoral Fellowship and an ESRC FRL Fellowship (ES/N017560/1) to SW.

